# Construction and optimization of multi-platform precision pathways for precision medicine

**DOI:** 10.1101/2023.05.23.541873

**Authors:** Andy Tran, Andy Wang, Jamie Mickaill, Dario Strbenac, Mark Larance, Steve Vernon, Stuart Grieve, Gemma Figtree, Ellis Patrick, Jean Yee Hwa Yang

**Affiliations:** School of Mathematics and Statistics, The University of Sydney, NSW, Australia; Charles Perkins Centre, The University of Sydney, NSW, Australia; Sydney Precision Data Science Centre, The University of Sydney, NSW, Australia; Sydney Medical School (Northern), The University of Sydney, NSW, Australia; School of Computer Science, The University of Sydney, NSW, Australia; Kolling Institute of Medical Research, St Leonards, NSW, Australia; Laboratory of Data Discovery for Health Limited (D^2^4H), Science Park, Hong Kong SAR, China

**Keywords:** translational research, multiomics, health economics

## Abstract

In the enduring challenge against disease, advancements in medical technology have empowered clinicians with novel diagnostic platforms. Whilst in some cases, a single test may provide a confident diagnosis, often additional tests are required. However, to strike a balance between diagnostic accuracy and cost-effectiveness, one must rigorously construct the clinical pathways. Here, we developed a framework to build multi-platform precision pathways in an automated, unbiased way, recommending the key steps a clinician would take to reach a diagnosis. We achieve this by developing a confidence score, used to simulate a clinical scenario, where at each stage, either a confident diagnosis is made, or another test is performed. Our framework provides a range of tools to interpret, visualize and compare the pathways, improving communication and enabling their evaluation on accuracy and cost, specific to different contexts. This framework will guide the development of novel diagnostic pathways for different diseases, accelerating the implementation of precision medicine into clinical practice.

## Introduction

In recent years, the medical field has seen rapid developments in various high-throughput biotechnologies, allowing the collection of biological data on a variety of “omics” platforms, at an increasingly scalable and affordable level^1^. For example, the cost of whole exome sequencing for a single sample has dramatically declined over the last two decades from around $20 million (USD) in 2006 to around $1000 (USD) in 2018^2^. This new access to a plethora of information is leading a revolution in precision medicine, by providing an insight into the biological mechanisms behind different diseases. Indeed, we already see modern omics data being used to help personalize cancer treatments^3^, among other diseases^4^. However, uptake of these technologies in clinical practice has been slow, due to the range of stakeholders involved^5^.

With so many novel technologies as potential diagnostic platforms^6^, much of current research aims to build a model for a cohort of patients using a single platform in isolation^7–9^, or integratively with other data^10,11^. However, this is different from the reality of a clinical application, where a range of diagnostic platforms/tests are available, and a variety of other factors need to be considered, such as health economics and time. In particular, with highly heterogeneous cohorts, a clinician may not necessarily need or want to perform such a test on all their patients, as there may be cheaper or more effective alternatives for some patients. Some recent research has aimed to identify clinical features to make cost-effective diagnosis under time and resource constraints^12^. Nonetheless for complex diseases, the specific order of testing, evaluation and clinical decision making is an important consideration, along with approaches for the integration of clinical, imaging and omics data^13^. For instance, genomic testing is rapidly transitioning into a “standard of care” to guide treatment plans for rare childhood diseases^14,15^ and cancer^16,17^.

Here, we present a framework (MultiP) to construct a multi-platform precision pathway given a range of available platforms to diagnose a disease (Figure 1). This workflow incorporates three main innovations: (1) An individualized confidence score to guide sequential decision-making for individuals, (2) the joint optimization of individual-level accuracy and population-level cost, and (3) interpretation tools to ensure a transparent and reliable implementation. We demonstrate the utility of our workflow in clinical and omics-level data for two different complex diseases (coronary artery disease) and stage III melanoma), outperforming a complete omics phenotyping, with less than half the number of diagnostic tests required. The framework is implemented in the R package ClassifyR, available on Bioconductor.

**Figure 1:**
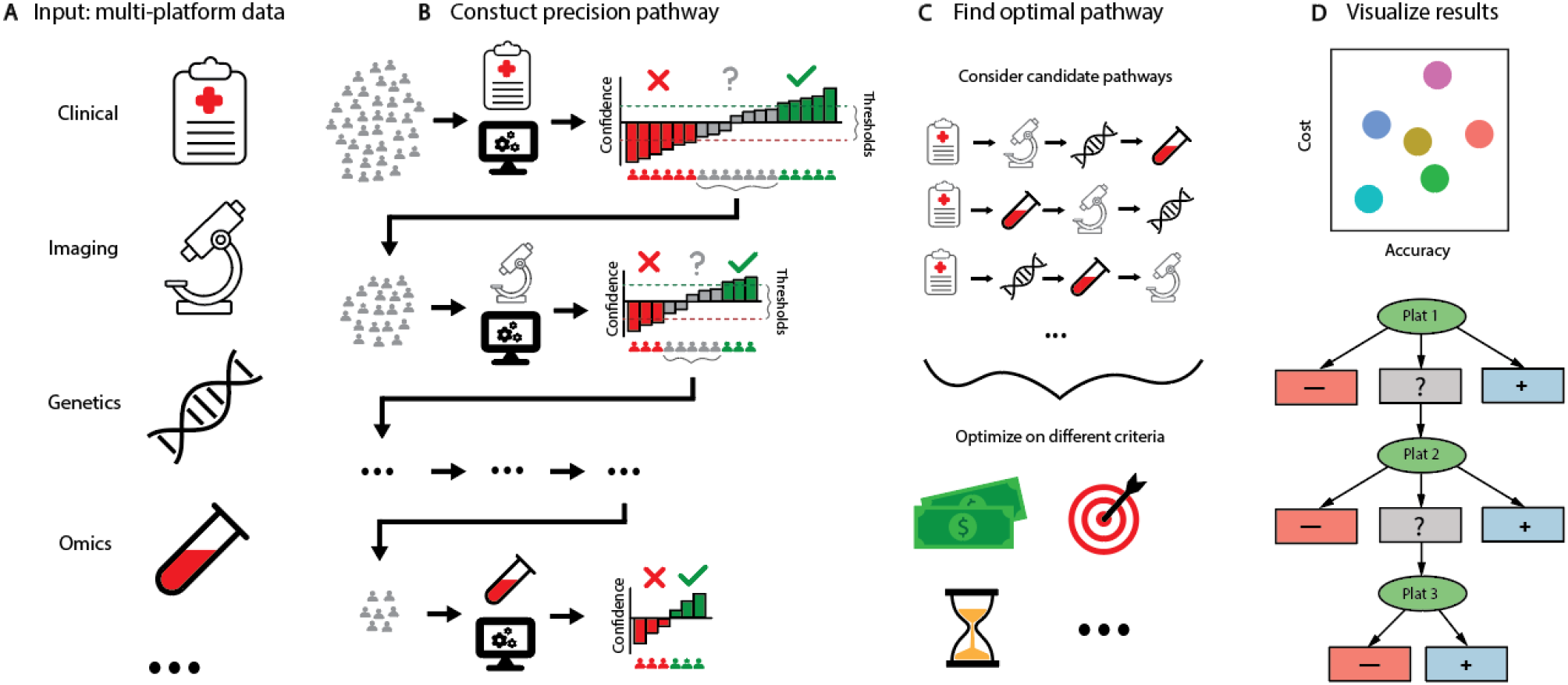
Schematic of the MultiP pipeline. **A**. The input into the model is data from multiple platforms for the same cohort of patients. **B**. For a particular sequence of platforms, the MultiP algorithm uses machine learning to classify patients into a positive, negative, or uncertain class. Patients in the uncertain class and passed onto the next platform and the process repeats until the final platform. **C**. Candidate pathways corresponding to different orders of platforms can be compared and optimized on different criteria. **D**. A suite of tools for visualizations and summaries of the constructed pathways are provided.

## Results

### The development of an “uncertain” class enables multi-stage classification

In a diagnostic setting, typical machine learning classifiers aim to classify all patients into either the positive or negative class^18^, which is a limited representation of the reality in the clinic. When given the results of a diagnostic test, a clinician may be able to make a diagnosis with high confidence, either positive or negative, or if they are “not sure”, they may refer the patient to take further tests to obtain more data. To capture this aspect of decision making, we introduce an “uncertain” class to the typical binary classification problem (turning it into a ternary classification problem), which allows us to perform a multi-stage classification, using the multiple modes of data available to us.

To build this multi-stage classification, we develop a confidence score for each patient and platform combination, representing the confidence in which we can make a diagnosis for that patient using that platform. We achieve this in a similar way to our previous work by Patrick and colleagues^19^, where a patient-specific accuracy rate is calculated by aggregating the predictions at a patient-level in repeated cross-validation. This can be imagined as having many different clinicians (models from each repeat in the cross-validation) to diagnose a new patient, and the confidence score is equivalent to the degree of agreement among the different clinicians. See *Methods (MultiP Algorithm: Confidence Score)* for full details.

We allow the user to customize the confidence threshold, as different contexts may require different stringencies. Based on this threshold, individuals can either be classified (if the model can make a high-confidence decision) or progressed (if the model cannot make a high-confidence decision). In the latter case, the individual proceeds to the next stage, where data from another platform will be collected. This process repeats until the final platform (all possible tests have been performed), where a decision must be made. See *Methods (MultiP Algorithm: Construction)* for full details.

Our MultiP framework is implemented as a part of the ClassifyR package^20^, available on Bioconductor. ClassifyR formalizes a framework for performing and evaluating classification in R using repeated cross-validation and includes many in-built feature selection and classification approaches. Many components of the ClassifyR framework can be customized with a single line of code, including the feature selection, classifier model and cross validation parameters making it ideal for implementing MultiP. This flexibility provides the versatility needed to cater to the varied contexts of different diseases and populations. The full list of parameters that can be adjusted in the MultiP framework are summarized in Table 1. See *Methods (Implementation)* for full details.

**Table 1:** Summary of parameters for MultiP

| Class | Parameter | Input | Default |
| --- | --- | --- | --- |
| MultiP parameters | Confidence Threshold | A value between 0 and 1 | 0.8 |
|  | Fixed Tiers | An integer from 0 to the number of platforms | 1 (corresponding to clinical data) |
|  | Platform Costs | Numerical (one for each platform) | NA |
|  | Criteria Weights | Numerical (one for each criteria) | (0.5, 0.5) |
|  | Additional Criteria Cost (optional) | Numerical (one for each platform) | NA |
| Feature Selection | Method | A character for a feature selection method | t-test |
|  | Number | A single number, or a vector of numbers (in which case, the optimal number from the vector will be selected) | (10, 20, 30, 40, 50, 60, 70, 80, 90, 100) |
| Classifier | Method | A character for a classifier | DLDA |
| Cross Validation | Cross-validation folds | Any positive integer | 2 |
|  | Repeats | Any positive integer | 50 |
|  | Workers | Any positive integer | 1 |
|  | Seed | Any positive integer | 1 |

### Clinical precision pathways made transparent by MultiP

When applying a machine learning model that can impact treatment decisions and people*’*s lives as well as the value and cost to the health system, it is imperative that the model is not treated as a black box, to ensure a robust and equitable implementation^21,22^. In MultiP, we ensure that the constructed pathways are interpretable, by providing a range of tools and visualizations to dissect the biomarkers driving the models. We demonstrate the utility and interpretability of MultiP on its ability to detect Coronary Artery Disease (CAD) in the BioHEART-CT cohort, using four platforms: clinical, metabolomics, lipidomics and proteomics. For full details about the cohort and data, see *Methods (Datasets)*.

For a single pathway, the flow chart provides an overall visualization of the progression of individuals at the population-level (Figure 2A). In our example, it can be seen that 78% of individuals can be confidently classified with just clinical information including standard modifiable risk factors, meaning that these individuals would not need any of the more expensive data to be collected. The major clinical unmet need is evident, where individuals without standard risk factors are in the subclinical phase of atherosclerotic development and at risk of heart attack, not detectable by traditional approaches. And equally, some individuals may appear to be at high risk based on clinical data alone (such as high cholesterol or smoking), but have distinct resilience, without the development of CAD. Knowledge of the latter may allow for avoidance of life-long pharmacotherapy.

**Figure 2:**
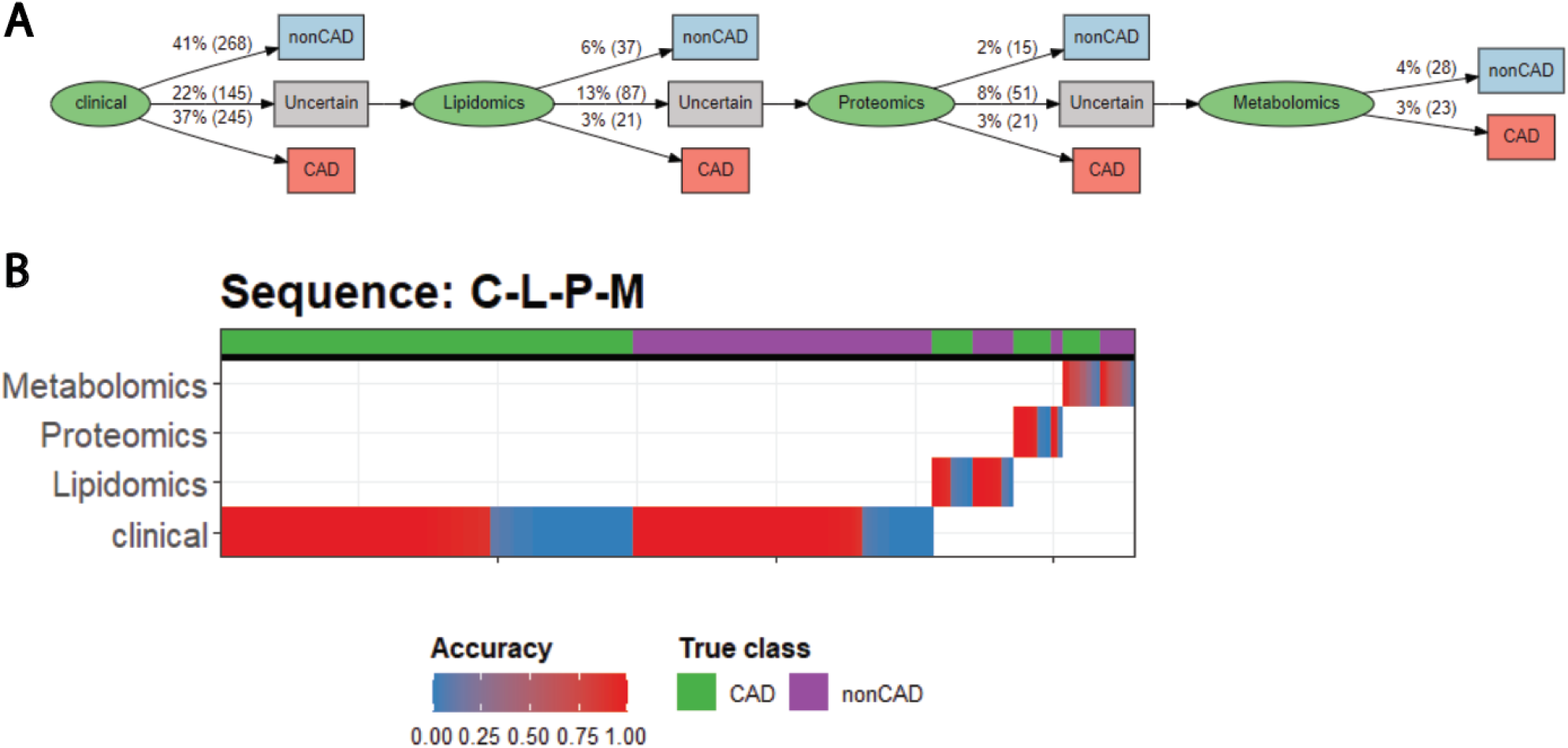
Visualization tools to interpret constructed pathways. **A**. Flow chart displays the proportion and number of patients assigned to each class at each stage of the pathway. **B**. Strata plot displays the accuracy of the patients classified in each stage of the pathway. The x-axis corresponds to each patient, sorted by the platform they were classified in (y-axis), then by their true class (top row), then by accuracy (color).

A more detailed look at the population can be seen in our strata plot that displays this data at the individual sample level (Figure 2B). At each stage of the pathway, the individuals are split by their true class, and the accuracy of the classifier is plotted for each individual. This can allow the user to assess the performance of each stage of the pathway and identify potential cohort heterogeneity. For instance, we see that the Lipidomics model of our MultiP has a high accuracy for non-CAD individuals, but low accuracy for CAD individuals. This suggests that this platform may have a bias for categorizing individuals as non-CAD, which could be investigated further.

To dissect the models created for each stage and identify the biomarkers that are driving the decision-making process, we produce a feature importance plot as shown in Figure S1A-C and Figure S2. Reassuringly, we see that in each of the models, the key features all have well-established links to CAD. Namely, the Lipidomics model (Figure S1A) is driven by Cer^23^, Hydroxylated acylcarnitine^24^, Sulfatide^25^; the Metabolomics model (Figure S1B) is driven by Riboflavin^26^, 2-arachidonoylglycerol^27^ and DMGV^28^; and the Proteomics model (Figure S1C) is driven by PON1^29^, IGFALS^30^ and SERPINC1^31^.

To examine the characteristics between classified and progressed individuals, we also provide a cohort summary of the classified and retained group of individuals at each stage of the pathway, to explore the differences in individual cohorts that are classified or progressed at each stage, potentially revealing cohort heterogeneity (Figure S3A-C). For instance, we see that the lipidomics model (Figure S3B) confidently classifies considerably more females than males, indicating a potential sex-bias in the data and/or model. This suggests that the different subpopulations (eg. sex) may be more appropriately classified with separate models, as has been well-studied in the context of CAD^32^.

### Criterion-guided optimization of precision pathways outperforms baseline models

To implement a multi-platform framework, an important consideration would be the “order of the platforms”, to represent the order that clinicians would perform diagnostic tests for individuals. In our MultiP framework, we train a precision pathway by first constructing a series of possible precision pathways for all possible orders of the platforms. Here the users can specify if any platforms must be used first, such as clinical data. Secondly, the final precision pathway is selected by comparing and assessing them based on accuracy at the population level.

To assess the performance of our constructed precision pathways, we compare their balanced accuracy to a range of baseline models. Here we consider two types of baseline models, the first group are models built on a single omics platform with clinical data, and the second group considers a model built on all platforms combined (Figure 3), see *Methods (Baseline comparison)* for full details about the construction of baseline models. We see that the precision pathways perform considerably better than single platform classifications, confirming the common belief that these different platforms contain complementary information, and demonstrating the value of integrating different platforms to make clinical diagnoses.

**Figure 3:**
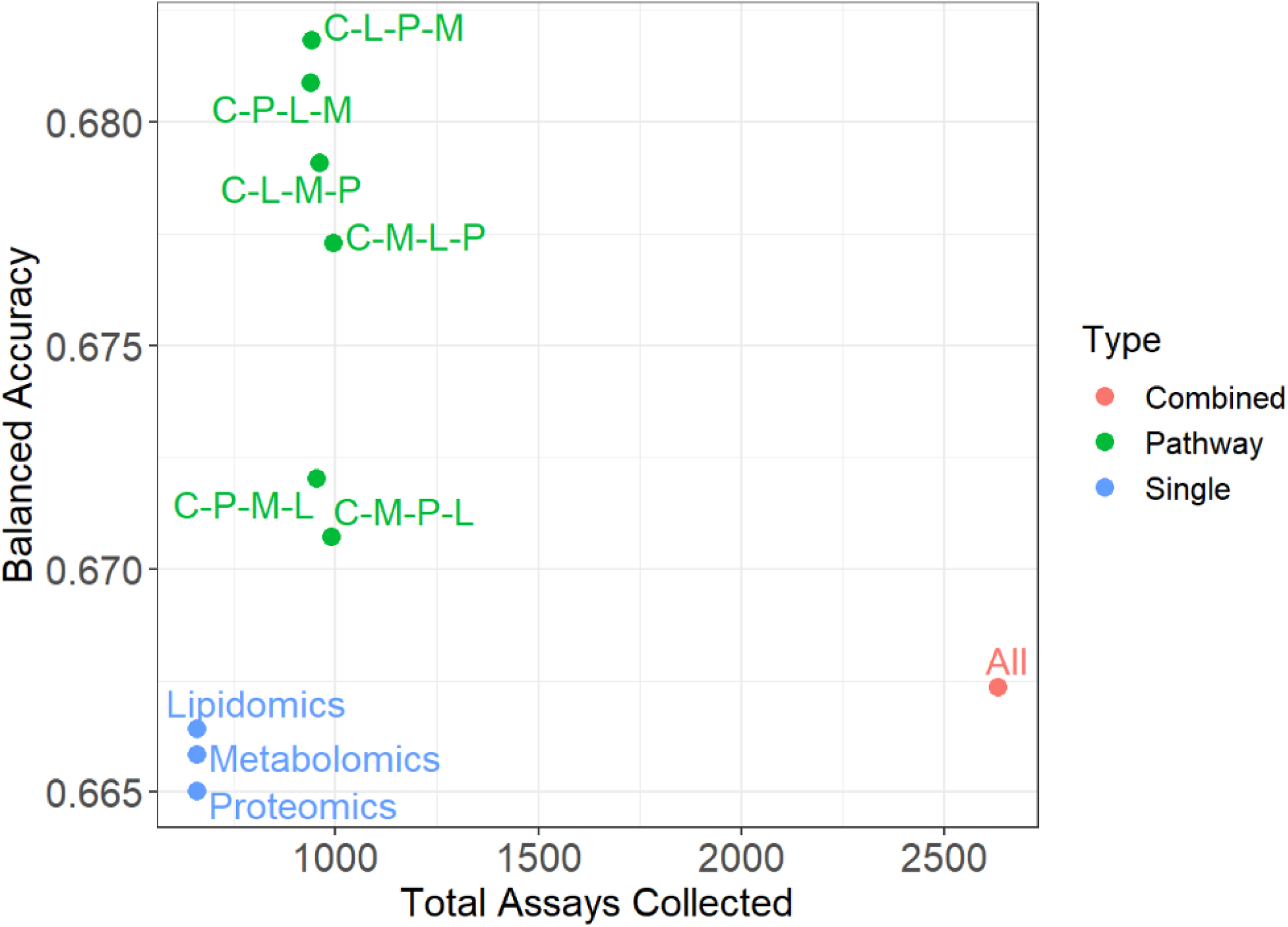
Comparison of precision pathways to baseline models. The x-axis corresponds to the total number of diagnostic assays that would need to be collected to diagnose the entire cohort and the y-axis corresponds to the repeated cross-validation balanced accuracy. The red point represents the model constructed on the full set of data; the green points represent the pathways built with MultiP; the blue points represent the models constructed on a single platform with clinical data.

The precision pathways not only maintain a high accuracy compared to the combined data set (as they are closer to the top of the plot), but only use a fraction of the data and thus a fraction of the cost (as they are closer to the left of the plot). This suggests that the combined data set, with all platforms for all patients, contains a large proportion of redundant information, i.e. there are many patients that can be confidently classified with little data. This highlights the value of a precision pathway to use only the important tests to reach confident diagnoses, while minimizing the cost of healthcare with limited sacrifice in accuracy.

### MultiP pathways can be optimized on multiple criteria

Classical machine learning approaches typically optimize their models based on accuracy alone^18^. However for a clinical implementation, there are a range of practical factors that need to be considered as well, such as cost or time. The MultiP framework allows users to incorporate additional criteria, and choose a weighting for the importance of each criterion when determining an optimal precision pathway. See *Methods (MultiP Algorithm: Evaluation)* for full details. A variety of tools and visualizations are provided to assist the user to compare the candidate models under these criteria.

Users can view a summary table of the constructed precision pathways, with their accuracy and cost at each level (Figure 4A). They are also given an overall score, based on their rankings for each of the criteria, aggregated by the user-chosen weightings. This allows for an unbiased selection of an optimal precision pathway. A bubble plot is also produced which allows users to compare the candidate precision pathways based on accuracy and cost (Figure 4B). Here, the ideal precision pathway would have a high accuracy and low cost and we see that there are three well-performing and economical pathways: C-L-P-M, C-L-M-P and C-M-L-P (where C = Clinical, L = Lipidomics, M = Metabolomics, P = Proteomics). However, the final choice of optimal pathway is a tradeoff between accuracy and cost.

**Figure 4:**
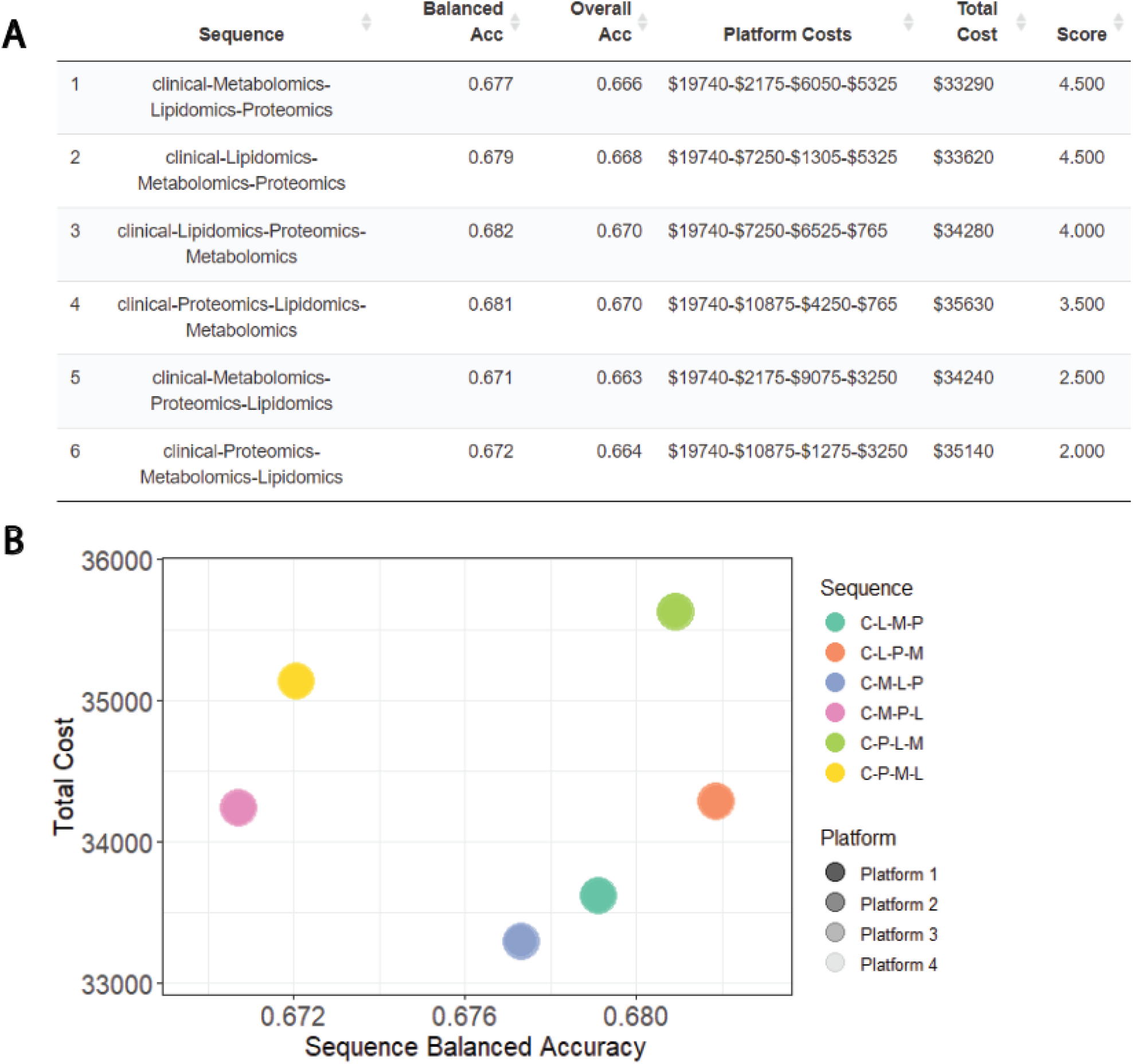
Visualization tools to compare candidate pathways. **A**. Summary table summarizes the accuracy and cost of each pathway. The pathways are ranked by score, calculated based on the pathways*’* rankings in balanced accuracy and cost, aggregated by user-defined weights. **B**. Bubble plot enables a visual comparison of the accuracy and cost of the candidate pathways. The shading of each point corresponds to the proportion of patients classified in each tier. Sequence names: C, clinical; L, lipidomics; M, metabolomics; P, proteomics.

### MultiP pathways are transferable across different cohorts

A major barrier to the implementation of new protocols for diagnostic purposes, such as omics technologies, is that molecular signatures are often cohort-specific and do not transfer well between different populations^33^. We further demonstrate the applicability and transferability of our framework for another complex disease, the prognosis of stage III melanoma. We achieve this by using MultiP to construct a pathway to classify patients into a “good prognosis” (survival > 4 years) or “poor prognosis” (survival < 1 year), trained on data from The Cancer Genome Atlas (TCGA)^34^. We then assess the performance of this model on a different dataset generated by the Melanoma Institute of Australia (MIA)^35–37^. See *Methods (Datasets)* for details about the cohorts.

Between these two cohorts, the data corresponding to the same molecular modality is sometimes generated from different technology platforms. In this situation, the microRNA data is generated using a count-based RNA-seq in the TCGA dataset and using a fluorescence-based microarray in the MIA dataset. To ensure transferability between the models, we use the log ratios between pairs of features as the input into the MultiP framework, as this has been demonstrated to be more appropriate for transferability^37^. See *Methods (Transferability analysis)* for full details.

We find that the models at each level are driven by well-known markers, suggesting that the constructed precision pathway is reasonable. In particular, we see that in the mRNA-level data (Figure S4B), the prediction for a poor prognosis is driven by higher levels of CCL21 and HAMP, both previously linked to the metastasis of melanomas^38,39^. The prediction for a good prognosis is driven by higher levels of DNAH2, a known modulator of cell homologous recombination repair which may have a protective effect^40^. In the microRNA model (Figure S4C), we observe that predictions for poor prognosis are driven by hsa-miR-205 and hsa-miR-518b, both known to be dysregulated in melanomas^41,42^. And a good prognosis is driven by hsa-miR-944 and hsa-miR-487a, known suppressors of cancer promoting genes^43,44^.

We then apply the precision pathway trained on the TCGA cohort to classify the individuals in the external MIA cohort (Figure 5). We find that the overall precision pathway maintains a good performance in balanced accuracy and F1 score (Table 2). However, we note that between the two cohorts, there was a considerable tradeoff between sensitivity and specificity. This is likely due to the small sample size available (65 in TCGA and 30 in MIA) resulting in overfitting to some subpopulations in the data.

**Figure 5:**
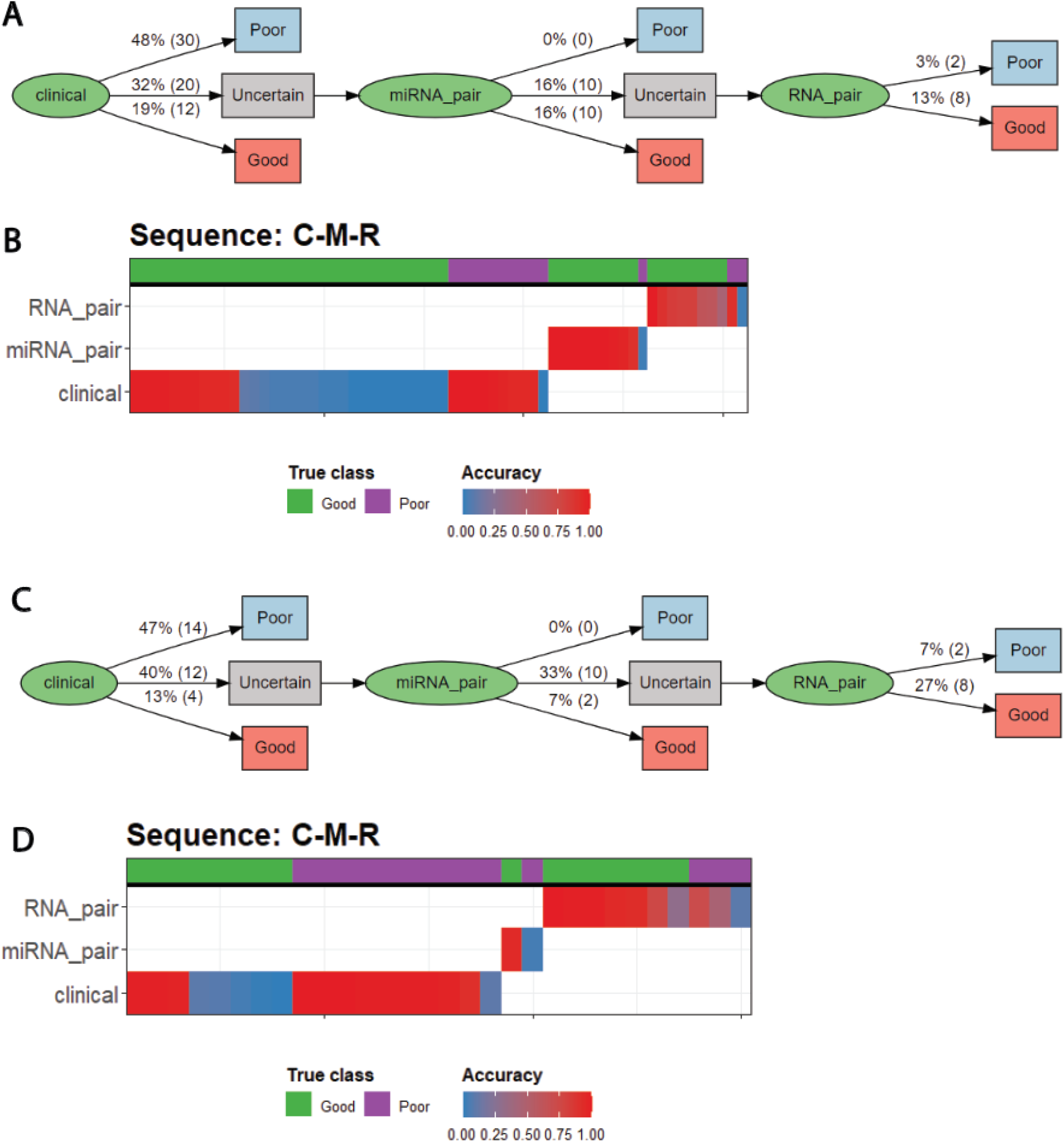
Demonstration of the transferability of MultiP. **A**. Flow chart for pathway on TCGA data. **B**. Strata plot for pathway on TCGA data. **C**. Flow chart for pathway on MIA data. **D** Strata plot for pathway on MIA data.

**Table 2:** Summary of performance metrics for model trained on TCGA dataset evaluated on TCGA dataset (with repeated cross-validation) and MIA dataset.

| Metric | TCGA Dataset | MIA Dataset |
| --- | --- | --- |
| Accuracy | 0.597 | 0.667 |
| Balanced Accuracy | 0.660 | 0.670 |
| F1 Score | 0.684 | 0.667 |
| Specificity | 0.769 | 0.714 |
| Sensitivity/Recall | 0.551 | 0.625 |
| Precision | 0.900 | 0.714 |

## Discussion

The increasing number of diagnostic platforms carry tremendous potential for clinical applications, but this brings the challenge of how to optimally use such data. Here, we present MultiP, a versatile framework to automatically construct precision pathways for a variety of contexts, using a range of available platforms. We provide a range of tools and visualizations to allow users to interpret the constructed pathways, and compare candidate pathways on different criteria. We have demonstrated the applicability of the framework in two distinct contexts: the diagnosis of CAD, and the prognosis of stage III melanoma on different cohorts.

We observed that these precision pathways have a similar performance to models constructed on a complete set of data, where data on all platforms are used for all patients. This implies that there are many patients for which a confident and accurate diagnosis can be made with minimal information, suggesting that it is not necessary to perform all tests for these patients. By following a precision pathway, we ensure that only the informative tests are performed, alleviating the huge economic burden of the healthcare system with minimal loss of accuracy.

The MultiP framework is implemented in the ClassifyR package, granting it access to a vast library of classification models and parameters in a single line of code. This flexibility allows the MultiP to cater to a wide range of contexts, as different diseases, populations and platforms would require tailored models. As further functionalities are incorporated into the ClassifyR package, such as new classifiers and multiview methods, MultiP will also expand in its applicability.

Despite the vast functionality and potential for MultiP to build diagnostic pathways, there remains a few limitations and scope for future work to improve its performance in a clinical application. Notably, cohort heterogeneity is a challenging issue, where there may be subpopulations in the cohort that would benefit from different classification models. The current MultiP uses the entire training cohort to build an ensemble model for all pathways, which may not accurately diagnose underrepresented subpopulations in the data. A workaround to this is to train a separate model at each stage of the pathway, so that the classifier used is more closely tailored to the subpopulation that is progressed, however these models will suffer from smaller sample size to train on. Another issue arising from cohort heterogeneity is that there may be different subpopulations that are better classified using a different order of platforms, whereas the current implementation forces the same order of platforms for all patient pathways. A solution could be that at each level, to identify potential subpopulations that are progressed and test different orders of platforms. However, this opens up exponentially more models that need to be constructed and tuned, quickly increasing the computational burden to construct the pathway.

In summary, our MultiP framework is the first to our knowledge that builds clinical pathways using multiple platforms and incorporating health economics. We hope that this will promote the uptake of modern technologies for clinical diagnoses, accelerating the progress towards precision medicine.

## Methods

### MultiP algorithm

#### Confidence score

The MultiP framework trains individual models for each platform using repeated cross-validation (with default parameters of 2 folds and 50 repeats). For an individual patient on a single platform, this will create many models (equal to the number of repeats) where the patient is in the test set, each with a predicted class. The final predicted class is then chosen based on the majority prediction across the many models. If the predictions are perfectly split across the two classes, the final predicted class is randomly chosen.

The confidence score is defined as the agreement of the predicted class of these models. More precisely, if the predicted classes are split in the ratio *p*: 1 − *p*, then the confidence score is defined as 2|*p* − 0.5|. That is, if all models predict the same class (*p* = 0 or *p* = 1), then the confidence score is 1, but if there is a perfect 50-50 split among the predictions (*p* = 0.5), then the confidence score is 0. For each patient in each platform, we have now calculated a final predicted class and a confidence score.

In our framework, we used default parameter values as stated in Table 1, and performed model building using the `runTests` function in ClassifyR^20^.

#### Construction

For a specific sequence of platforms and a user-defined confidence threshold, the pathway is constructed as follows:

1. All patients start at the first platform.
2. At the current platform, classify the patients, whose confidence score for that platform exceeds the threshold, with their final predicted class.
3. For the patients whose confidence score does not exceed the threshold, they are considered “uncertain” and then passed onto the next platform.
4. Repeat steps 2 and 3 until the final platform.
5. At the final platform, classify all patients based on the final predicted class for that platform.

We use a default confidence threshold of 0.8, however the optimal threshold may vary greatly across contexts, as different diseases, populations and platforms would have different accuracies and confidence.

#### Evaluation

When a pathway is constructed, each patient is assigned a predicted class. By comparing these predictions to their true class, we can calculate any classification metric, such as accuracy, balanced accuracy, F1 score, and so on.

To evaluate a list of candidate pathways, we assign a ranking to each one in each criteria, such as accuracy and cost. A weighted average of these rankings, based on user-defined weights, is taken to be the final score used to determine the optimal pathway. We choose default weights of 0.5 for accuracy and 0.5 for cost.

### Datasets

#### BioHEART-CT

This study has been described in detail previously^45^ and we analyze the Discovery 1000 patients, which is the first 1000 patients of the BioHEART-CT study who have completed deep imaging and molecular phenotyping. The study was approved by the Northern Sydney Local Health District Human Research Ethics Committee (HREC/17/HAWKE/343) and all participants provided informed written consent. All methods were performed in accordance with relevant guidelines and regulations. The deep imaging (CTCA images) were acquired on a 256-slice scanner using standard clinical protocols, overseen and dual-reported by accredited cardiologists and radiologists. CTCAs were analyzed using the validated 17-segment Gensini score^46^ to identify those with CAD (Gensini > 0) and without CAD (Gensini = 0). Data from molecular phenotyping include proteomics, lipidomics and metabolomics^47^. For demonstration purposes, the cost of each platform was chosen to be: Clinical = $30, Lipidomics = $50, Metabolomics = $15, Proteomics = $75. Patients on statin medications were excluded from analysis, as this would have an undesired confounding effect onto the molecular signatures.

#### The Cancer Genome Atlas (TCGA)

The SKCM (Skin Cutaneous Melanoma) data set was downloaded from TCGA using the R package`curatedTCGAData` ^48^.The `RNASeq2GeneNorm` and `miRNASeqGene` assays were taken to represent the mRNA and microRNA platforms respectively. The cohort was filtered down to those with stage III cancers to match with the MIA dataset. A “Good” prognosis was defined to be survival greater than 4 years from the date of tumor banking, and a “Poor” prognosis was defined to be death less than 1 year from the date of tumor banking. Patients who do not match a “Good” or “Poor” prognosis are excluded from analysis. The T-stages of the patients were reclassified into T0, T1, T2, T3 and T4, where patients with missing or undetermined T-stage were excluded.

#### Melanoma Institute of Australia (MIA)

This data collection includes data presented in Mann et al.^35^ and Jayawardana et al.^36^ and accessible at Melanoma Explorer^49^. In brief, mRNA was assayed using Sentrix Human-6 v3 Expression BeadChips (Illumina, San Diego, CA) and microRNA expression profiling was performed using Agilent Technologies*’* microRNA platform (version 16, Agilent Technologies, Santa Clara, CA). Similarly to the TCGA dataset, a “Good” prognosis was defined to be survival greater than 4 years from the date of tumor banking, and a “Poor” prognosis was defined to be death less than 1 year from the date of tumor banking. Patients who do not match a “Good” or “Poor” prognosis are excluded from analysis.

### Baseline comparison

To evaluate the performance of pathways generated by MultiP, we compare against two categories of models. Single platform models are built on each individual platform with clinical data and the combined model was built on all platforms integratively. The integration of different platforms for classification was implemented with the `crossValidate` function in ClassifyR with the parameter `multiViewMethod = “merge”`. The same cross-validation parameters as the MultiP pathways were used to ensure a fair comparison.

### Transferability analysis

To build a transferable model between the TCGA and MIA datasets, we first perform library size normalization and then filter the features in each platform to those that are common in both datasets. In the TCGA data (the training data), we filter the microRNA features to those with standard deviation greater than 5, to keep the number of features reasonable for the next step while retaining important features.

As the data was collected from different platforms, with values on different scales, we calculate the log-ratios between each pair of features, using the method described by Wang and colleagues^37^. We then standardize these log ratios at the patient-level, shifting the mean to 0 and scaling the variance to 1. Pairs with very low standard deviation (< 0.1) are removed from analysis to ensure model stability.

### Implementation

The MultiP framework is implemented in the ClassifyR package available on Github at https://github.com/SydneyBioX/ClassifyR. It will also be available on Bioconductor in the next release.

## Data and code availability

The TCGA data that support the findings of this study are publicly available at the Genomic Data Commons Data Portal (Project ID: TCGA-SKCM). For the BioHEART-CT data, data requests can be made through the BioHEART data committee via email. The code for running the above methods and evaluation are available at https://github.com/SydneyBioX/ClassifyR.

## Author Contributions

JY conceived the design of the project. AT led the computational development described in this work with contributions from AW, JM and DS, with supervision from EP and JY. GF and SG led the BioHEART-CT study with contributions from SV. ML led the proteomics component and SG led the imaging component of the BioHEART-CT study respectively.

## Acknowledgements

The authors thank the staff at the Royal North Shore Hospital, in particular staff at North Shore Radiology and staff at NSW Health Pathology for their support. The authors also thank their colleagues at the University of Sydney, School of Mathematics and Statistics, Charles Perkins Center, and Kolling Institute for their intellectual engagement.

