## Supplementary figures for "Construction and optimization of multi-platform precision pathways for precision medicine"

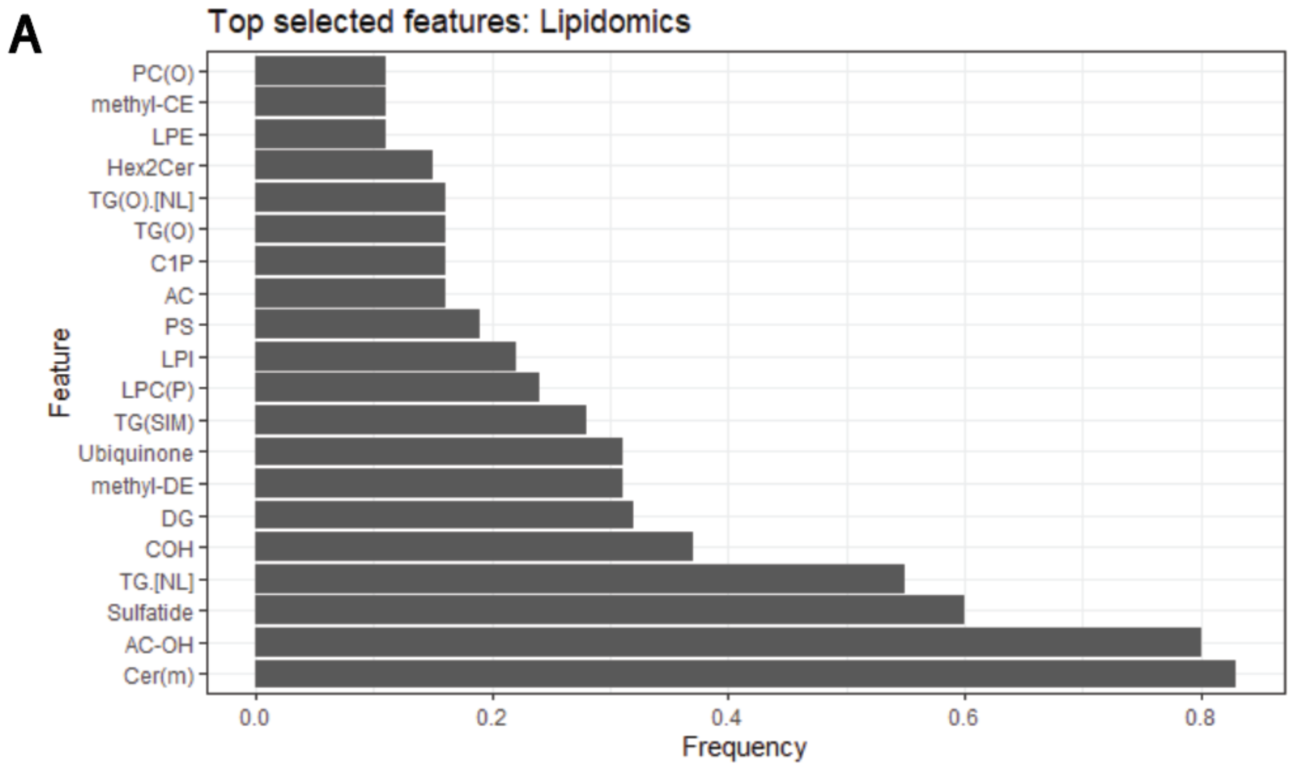


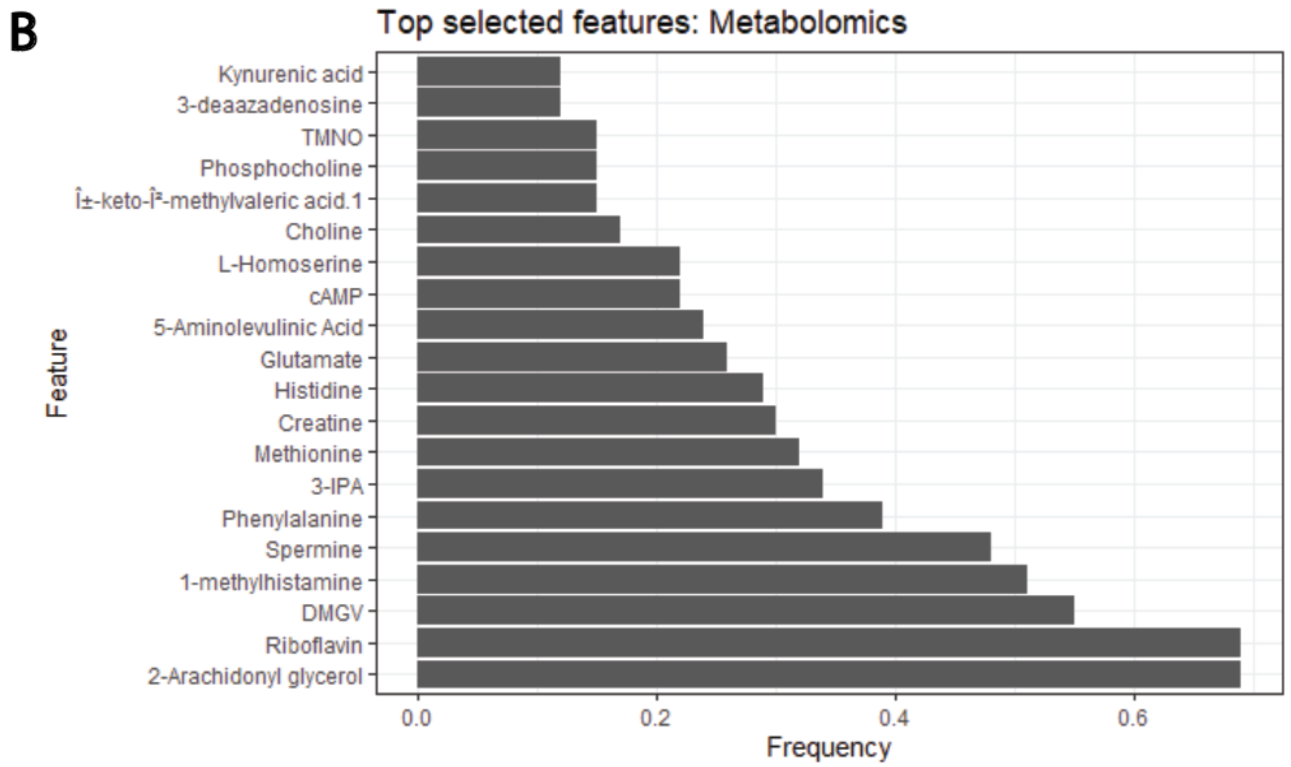

C

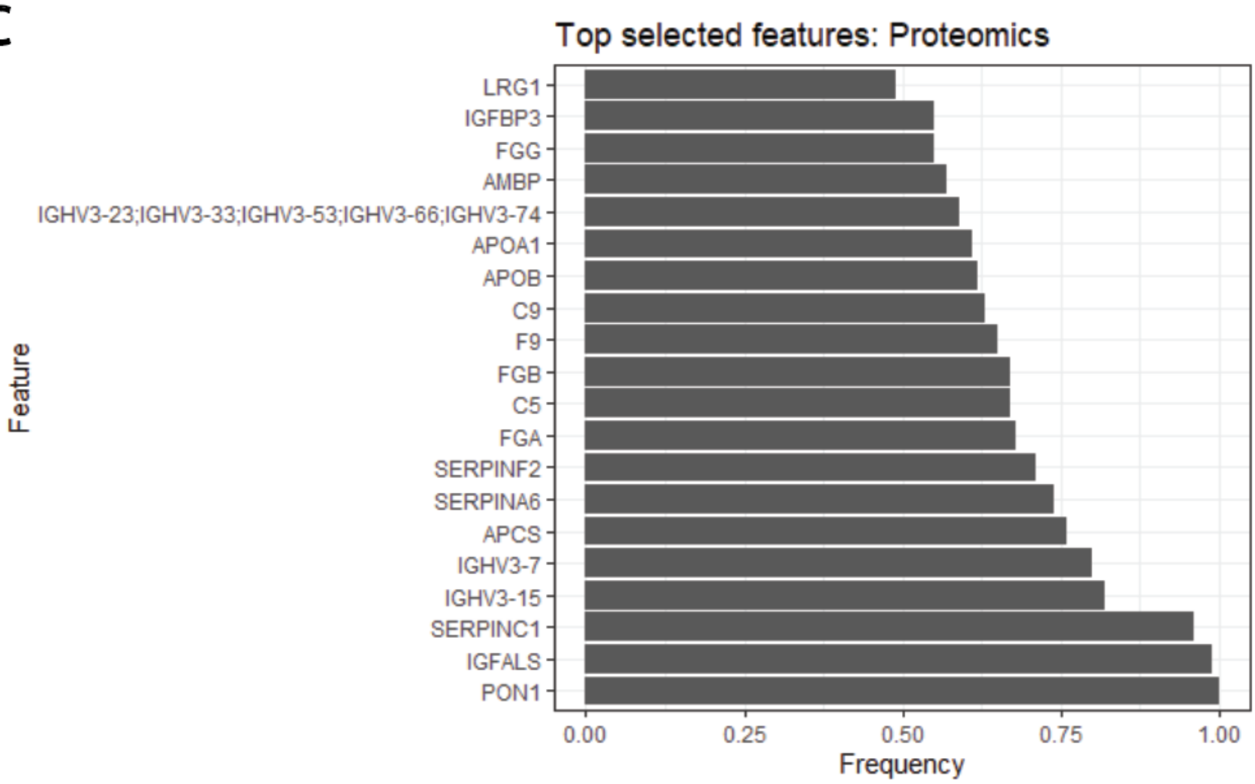


**Supplementary Figure 1:** Feature selection plots visualize the proportion of folds where each feature was selected, indicative of the importance of each feature to the overall model. **A**. Feature selection plot for Lipidomics in detecting CAD. **B**. Feature selection plot for Metabolomics in detecting CAD. **C**. Feature selection plot for Proteomics in detecting CAD.

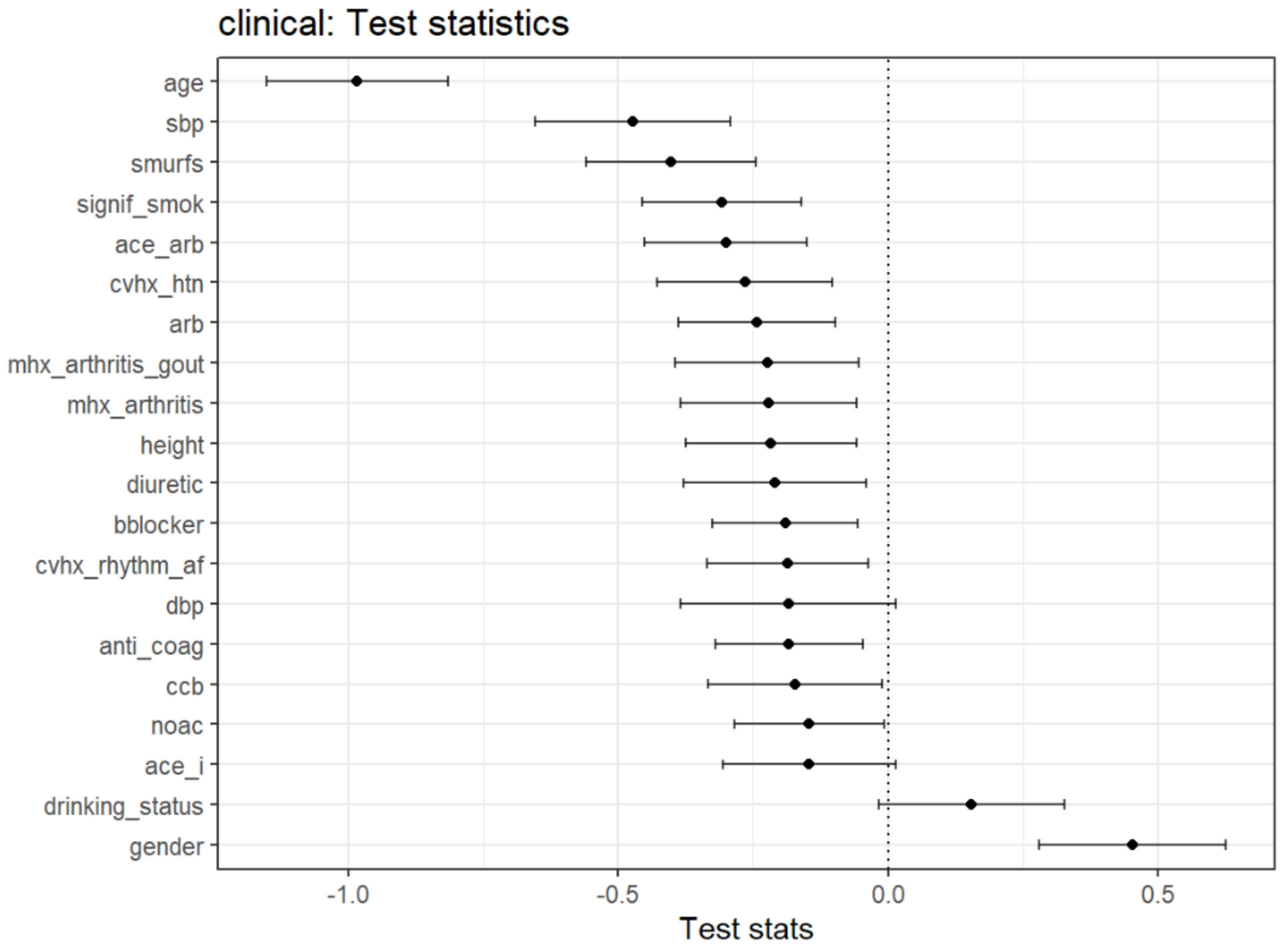


**Supplementary Figure 2:** DLDA feature importance plot visualizes the loadings of each feature in the discriminant function, indicative of the importance of each feature. Error bars correspond to a 95% confidence interval for the loadings, estimated from the repeated cross-validation. The above results plot the values for clinical data in detecting CAD. age = age in years, sbp = systolic blood pressure, smurfs = standard modifiable risk factors, signif_smok = smoking pack year history > 10 years, ace_arb = ACE inhibitor/ARB, cvhx_htn = hypertension, arb = ARB, mhx_arthiritis_gout = gout, mhx_arthiritis = osteoarthritis, height = height in meters, diuretic = diuretic, bblocker = beta-blocker, cvhx_rhythm_af = atrial fibrillation, dbp = diastolic blood pressure, anti_coag = anti-coagulant, ccb = calcium channel blocker, noac = NOAC, ace_i = ace inhibitor, drinking_status = drinking status (current, ex-drinker, never), gender = sex (male, female)

**A**

| clinical | Retained (N=513) | To progress (N=145) | Overall (N=658) |
|---|---|---|---|
| **height** | | | |
| Mean (SD) | 1.72 (0.104) | 1.72 (0.101) | 1.72 (0.103) |
| Median [Min, Max] | 1.71 [1.40, 2.00] | 1.72 [1.49, 1.97] | 1.71 [1.40, 2.00] |
| **smurfs** | | | |
| Mean (SD) | 0.924 (0.857) | 0.986 (0.645) | 0.938 (0.815) |
| Median [Min, Max] | 1.00 [0, 4.00] | 1.00 [0, 3.00] | 1.00 [0, 4.00] |
| **cvhx_htn** | | | |
| 0 | 348 (67.8%) | 101 (69.7%) | 449 (68.2%) |
| 1 | 165 (32.2%) | 44 (30.3%) | 209 (31.8%) |
| **cvhx_hcl_sr** | | | |
| 0 | 316 (61.6%) | 88 (60.7%) | 404 (61.4%) |
| 1 | 197 (38.4%) | 57 (39.3%) | 254 (38.6%) |
| **smoking_status** | | | |
| 1 | 30 (5.8%) | 15 (10.3%) | 45 (6.8%) |
| 2 | 293 (57.1%) | 77 (53.1%) | 370 (56.2%) |
| 3 | 190 (37.0%) | 53 (36.6%) | 243 (36.9%) |
| **drinking_status** | | | |
| 1 | 401 (78.2%) | 116 (80.0%) | 517 (78.6%) |
| 2 | 13 (2.5%) | 5 (3.4%) | 18 (2.7%) |
| 3 | 89 (17.3%) | 21 (14.5%) | 110 (16.7%) |
| 4 | 10 (1.9%) | 3 (2.1%) | 13 (2.0%) |
| **sbp** | | | |
| Mean (SD) | 131 (17.4) | 131 (14.2) | 131 (16.8) |
| Median [Min, Max] | 129 [89.0, 197] | 130 [99.0, 173] | 130 [89.0, 197] |
| **dbp** | | | |
| Mean (SD) | 75.6 (10.6) | 76.5 (11.1) | 75.8 (10.7) |
| Median [Min, Max] | 76.0 [41.0, 106] | 77.0 [45.0, 111] | 76.0 [41.0, 111] |
| **hr** | | | |
| Mean (SD) | 66.8 (11.8) | 66.3 (10.9) | 66.7 (11.6) |
| Median [Min, Max] | 66.0 [35.0, 115] | 66.0 [35.0, 111] | 66.0 [35.0, 115] |
| **age** | | | |
| Mean (SD) | 58.2 (12.7) | 59.7 (10.3) | 58.6 (12.2) |
| Median [Min, Max] | 59.0 [21.0, 92.0] | 60.0 [27.0, 82.0] | 59.0 [21.0, 92.0] |
| **gender** | | | |
| 1 | 291 (56.7%) | 75 (51.7%) | 366 (55.6%) |
| 2 | 222 (43.3%) | 70 (48.3%) | 292 (44.4%) |

**B**

| Lipidomics | Retained (N=58) | To progress (N=87) | Overall (N=145) |
|---|---|---|---|
| **height** | | | |
| Mean (SD) | 1.70 (0.119) | 1.72 (0.0869) | 1.72 (0.101) |
| Median [Min, Max] | 1.70 [1.49, 1.97] | 1.72 [1.57, 1.94] | 1.72 [1.49, 1.97] |
| **smurfs** | | | |
| Mean (SD) | 1.10 (0.612) | 0.908 (0.658) | 0.986 (0.645) |
| Median [Min, Max] | 1.00 [0, 3.00] | 1.00 [0, 2.00] | 1.00 [0, 3.00] |
| **cvhx_htn** | | | |
| 0 | 33 (56.9%) | 68 (78.2%) | 101 (69.7%) |
| 1 | 25 (43.1%) | 19 (21.8%) | 44 (30.3%) |
| **cvhx_hcl_sr** | | | |
| 0 | 34 (58.6%) | 54 (62.1%) | 88 (60.7%) |
| 1 | 24 (41.4%) | 33 (37.9%) | 57 (39.3%) |
| **smoking_status** | | | |
| 1 | 10 (17.2%) | 5 (5.7%) | 15 (10.3%) |
| 2 | 30 (51.7%) | 47 (54.0%) | 77 (53.1%) |
| 3 | 18 (31.0%) | 35 (40.2%) | 53 (36.6%) |
| **drinking_status** | | | |
| 1 | 48 (82.8%) | 68 (78.2%) | 116 (80.0%) |
| 2 | 1 (1.7%) | 4 (4.6%) | 5 (3.4%) |
| 3 | 8 (13.8%) | 13 (14.9%) | 21 (14.5%) |
| 4 | 1 (1.7%) | 2 (2.3%) | 3 (2.1%) |
| **sbp** | | | |
| Mean (SD) | 131 (14.6) | 130 (14.0) | 131 (14.2) |
| Median [Min, Max] | 129 [99.0, 171] | 132 [101, 173] | 130 [99.0, 173] |
| **dbp** | | | |
| Mean (SD) | 76.7 (10.6) | 76.4 (11.4) | 76.5 (11.1) |
| Median [Min, Max] | 76.0 [56.0, 111] | 77.0 [45.0, 99.0] | 77.0 [45.0, 111] |
| **hr** | | | |
| Mean (SD) | 66.2 (11.5) | 66.3 (10.5) | 66.3 (10.9) |
| Median [Min, Max] | 66.0 [43.0, 111] | 65.0 [35.0, 89.0] | 66.0 [35.0, 111] |
| **age** | | | |
| Mean (SD) | 58.8 (9.70) | 60.3 (10.7) | 59.7 (10.3) |
| Median [Min, Max] | 60.0 [27.0, 80.0] | 60.0 [29.0, 82.0] | 60.0 [27.0, 82.0] |
| **gender** | | | |
| 1 | 26 (44.8%) | 49 (56.3%) | 75 (51.7%) |
| 2 | 32 (55.2%) | 38 (43.7%) | 70 (48.3%) |

C

| Proteomics | Retained (N=36) | To progress (N=51) | Overall (N=87) |
|---|---|---|---|
| **height** | | | |
| Mean (SD) | 1.72 (0.0762) | 1.73 (0.0944) | 1.72 (0.0869) |
| Median [Min, Max] | 1.71 [1.57, 1.91] | 1.73 [1.57, 1.94] | 1.72 [1.57, 1.94] |
| **smurfs** | | | |
| Mean (SD) | 1.00 (0.676) | 0.843 (0.644) | 0.908 (0.658) |
| Median [Min, Max] | 1.00 [0, 2.00] | 1.00 [0, 2.00] | 1.00 [0, 2.00] |
| **cvhx_htn** | | | |
| 0 | 30 (83.3%) | 38 (74.5%) | 68 (78.2%) |
| 1 | 6 (16.7%) | 13 (25.5%) | 19 (21.8%) |
| **cvhx_hcl_sr** | | | |
| 0 | 22 (61.1%) | 32 (62.7%) | 54 (62.1%) |
| 1 | 14 (38.9%) | 19 (37.3%) | 33 (37.9%) |
| **smoking_status** | | | |
| 1 | 2 (5.6%) | 3 (5.9%) | 5 (5.7%) |
| 2 | 18 (50.0%) | 29 (56.9%) | 47 (54.0%) |
| 3 | 16 (44.4%) | 19 (37.3%) | 35 (40.2%) |
| **drinking_status** | | | |
| 1 | 25 (69.4%) | 43 (84.3%) | 68 (78.2%) |
| 2 | 3 (8.3%) | 1 (2.0%) | 4 (4.6%) |
| 3 | 8 (22.2%) | 5 (9.8%) | 13 (14.9%) |
| 4 | 0 (0%) | 2 (3.9%) | 2 (2.3%) |
| **sbp** | | | |
| Mean (SD) | 127 (14.8) | 133 (13.1) | 130 (14.0) |
| Median [Min, Max] | 127 [101, 154] | 132 [109, 173] | 132 [101, 173] |
| **dbp** | | | |
| Mean (SD) | 74.7 (12.2) | 77.5 (10.8) | 76.4 (11.4) |
| Median [Min, Max] | 75.5 [54.0, 99.0] | 78.0 [45.0, 95.0] | 77.0 [45.0, 99.0] |
| **hr** | | | |
| Mean (SD) | 66.7 (12.2) | 66.0 (9.35) | 66.3 (10.5) |
| Median [Min, Max] | 64.0 [35.0, 89.0] | 65.0 [49.0, 88.0] | 65.0 [35.0, 89.0] |
| **age** | | | |
| Mean (SD) | 62.0 (11.1) | 59.1 (10.4) | 60.3 (10.7) |
| Median [Min, Max] | 63.0 [41.0, 82.0] | 59.0 [29.0, 81.0] | 60.0 [29.0, 82.0] |
| **gender** | | | |
| 1 | 22 (61.1%) | 27 (52.9%) | 49 (56.3%) |
| 2 | 14 (38.9%) | 24 (47.1%) | 38 (43.7%) |

**Supplementary Figure 3:** Cohort summary tables summarize the cohorts that are classified or progressed at each stage of the pathway. **A**. Cohort summary table for stage 1 (clinical). **B**. Cohort summary table for stage 2 (lipidomics). **C**. Cohort summary table for stage 3 (metabolomics).

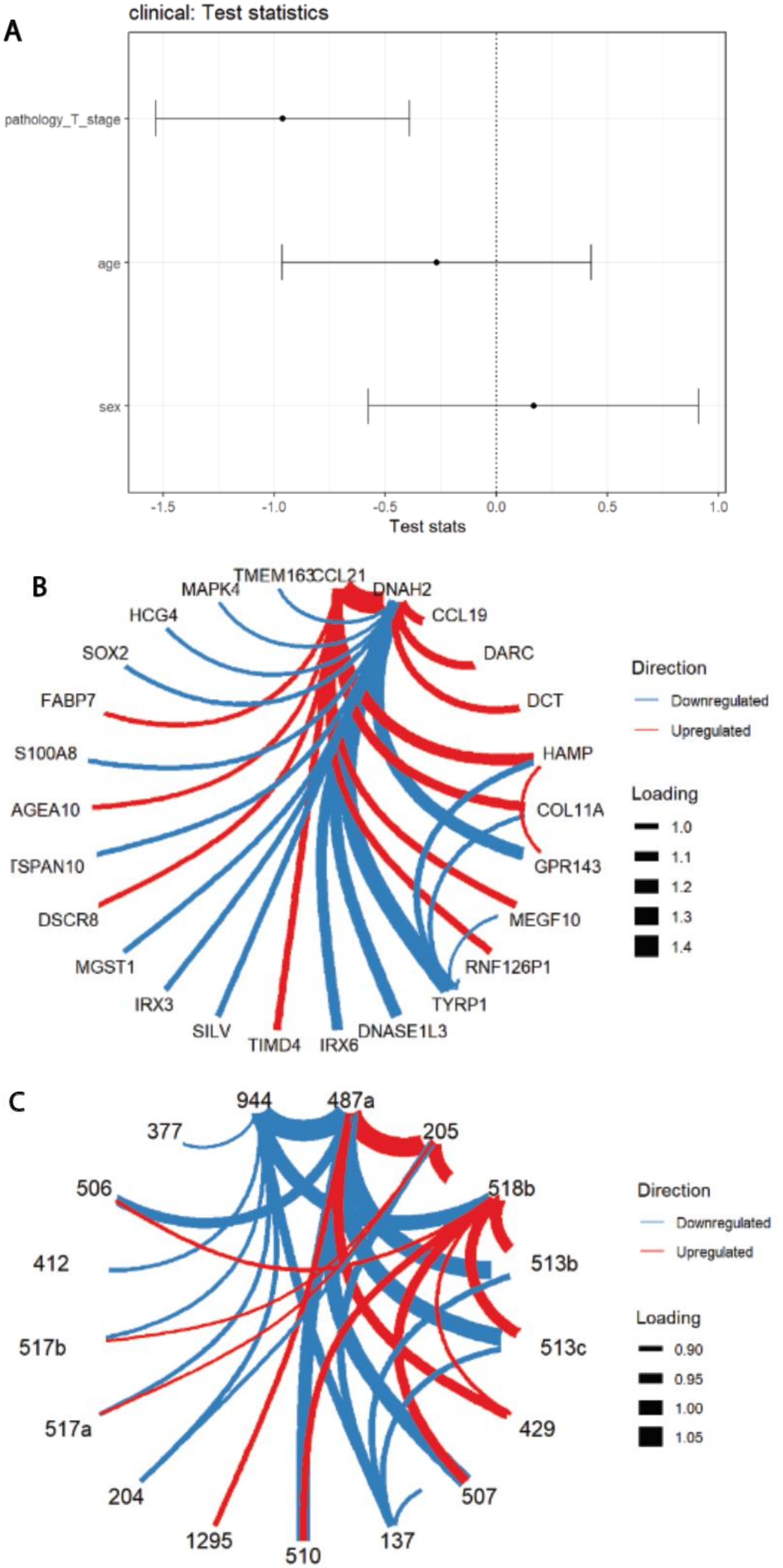

A
clinical: Test statistics
pathology_T_stage
age
sex
-1.5
-1.0
-0.5
0.0
0.5
1.0
Test stats
B
TMEM163
CCL21
DNAH2
MAPK4
HCG4
SOX2
FABP7
S100A8
AGEA10
TSPAN10
DSCR8
MGST1
IRX3
SILV
TIMD4
IRX6
DNASE1L3
TYRP1
RNF126P1
MEGF10
GPR143
COL11A
HAMP
DCT
DARC
CCL19
Direction
Downregulated
Upregulated
Loading
1.0
1.1
1.2
1.3
1.4
C
944
487a
377
205
506
518b
412
513b
517b
513c
517a
429
204
507
1295
510
137
Direction
Downregulated
Upregulated
Loading
0.90
0.95
1.00
1.05

**Supplementary Figure 4:** Feature importance plots for each model built on the TCGA dataset for melanoma prognosis. For models built on log ratios between pairs of features, the DLDA discriminant function feature loadings are visualized as a connection between pairs of features. The thickness of connection is proportional to the magnitude of feature loadings, and color corresponds to the sign of the feature loading. **A**. DLDA Feature importance plot of clinical data. **B**. DLDA Feature importance plot of mRNA data (log ratios). **C**. DLDA Feature importance plot of microRNA data (log ratios).
